# Neural Coding of Noisy and Reverberant Speech in Human Auditory Cortex

**DOI:** 10.1101/229153

**Authors:** Krishna C Puvvada, Marisel Villafañe-Delgado, Christian Brodbeck, Jonathan Z Simon

## Abstract

Speech communication in daily listening environments is complicated by the phenomenon of reverberation, wherein any sound reaching the ear is a mixture of the direct component from the source and multiple reflections off surrounding objects and the environment. The brain plays a central role in comprehending speech accompanied by such distortion, which, frequently, is further complicated by the presence of additional noise sources in the vicinity. Here, using magnetoencephalography (MEG) recordings from human subjects, we investigate the neural representation of speech in noisy, reverberant listening conditions as measured by phase-locked MEG responses to the slow temporal modulations of speech. Using systems-theoretic linear methods of stimulus encoding, we observe that the cortex maintains both distorted and distortion-free (cleaned) representations of speech. Also, we show that, while neural encoding of speech remains robust to additive noise in absence of reverberation, it is detrimentally affected by noise when present along with reverberation. Further, using linear methods of stimulus reconstruction, we show that theta-band neural responses are a likely candidate for the distortion free representation of speech, whereas delta band responses are more likely to carry non-speech specific information regarding the listening environment.

## Introduction

Speech communication in real-world scenarios, such as in a room or other enclosed space, differs from communication in an isolated environment as the sound entering the ear is a linear superposition of direct (clean, distortion-free) component and multiple reflections from the surroundings. This general acoustic phenomenon, known as reverberation, is ubiquitous in daily listening environments. The reflections travel a longer path, with correspondingly attenuated amplitudes, before summing linearly with the direct component, thus distorting the clean sound from the original source. Depending on the number of reflections and their attenuation factors (a function itself of the surrounding reflecting surfaces and the paths travelled), the distortion of clean sound can vary from mild (e.g., in large open spaces) to severe (e.g., in a cave, cathedral or a dense forest). The reverberant signal received by the ear can be modeled as *y*(*t*) = *s*(*t*)\**h*(*t*), where *s*(*t*) is the clean sound from the source and *h*(t) is the impulse response of a linear filter representing the delay and attenuation information of reflections (Figure 1**Error! Reference source not found**.). On the other hand, knowing only the reverberant signal *y*(*t*), to infer the original sound *s*(*t*) without knowledge of *h*(*t*) is mathematically ill-posed problem, though human listeners are nonetheless able to perform this routinely, with some effort (Sarampalis et al. 2009; Sato et al. 2007; Yang and Bradley 2009). Comprehension of speech in such a reverberant environment is further complicated by the presence of other sound sources whether stationary (e.g., the sound of an air-conditioner) or non-stationary (e.g., other talkers). The neural mechanisms by which reverberation is accommodated, and the representations employed by the auditory system in that process, in such adverse listening conditions remains unclear.

**Figure 1.**
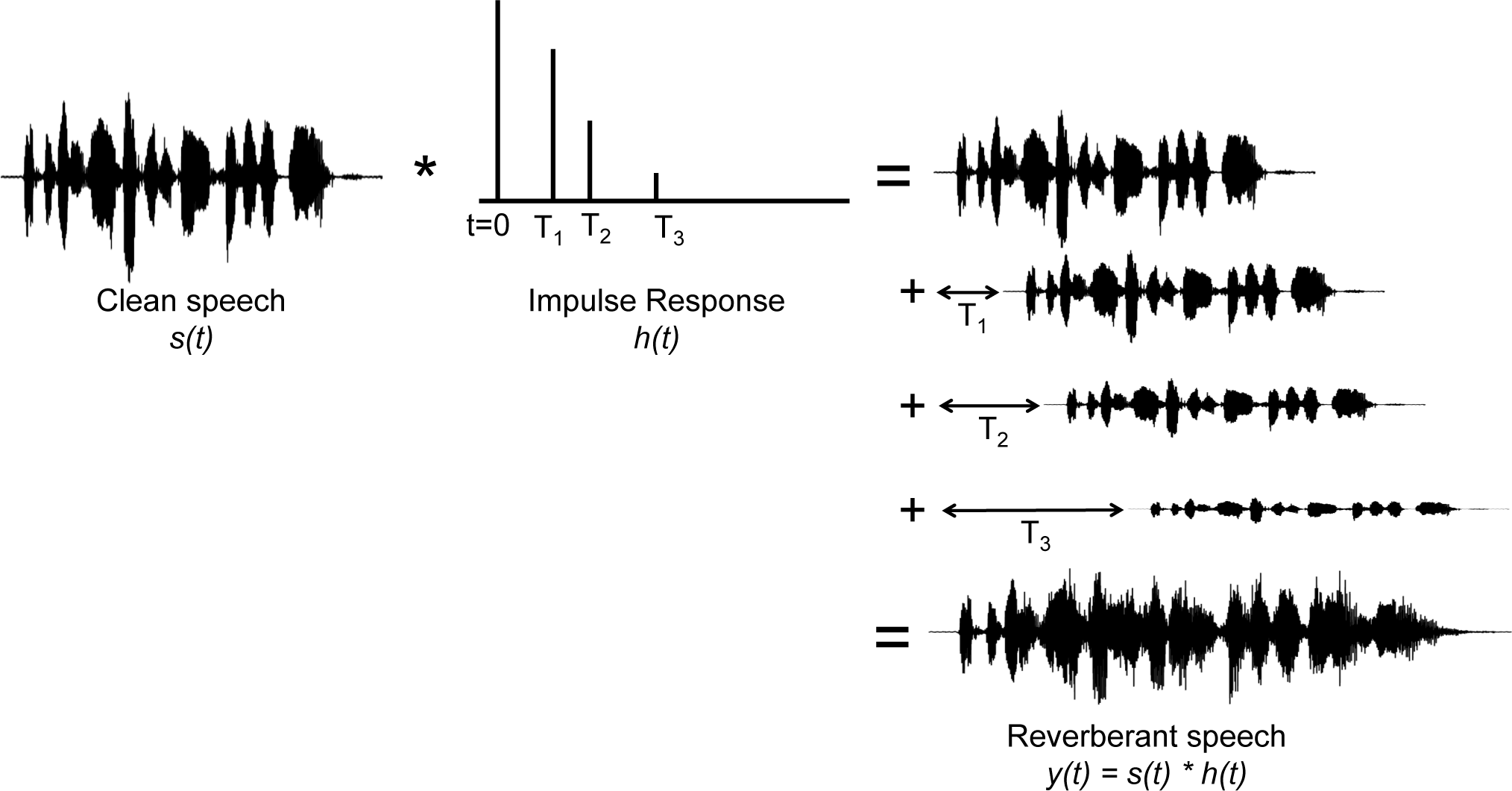
Phenomenon of reverberation. A reverberant signal reaching the ear is the sum of the original clean speech and its copies, appropriately time-shifted and scaled. This can be described mathematically as convolution between the clean speech *s*(*t*) and the reverberation impulse response *h*(*t*) (illustrated here with a schematic impulse response; after Traer and McDermott (Traer and McDermott 2016))

The information in speech is conveyed through its temporal modulations, which can be decomposed into a slow envelope that modulates the fast temporal fine structure (TFS) (Rosen 1992; Shamma and Lorenzi 2013). The slower envelope (<10 Hz) corresponds to prosodic, phonemic, syllabic and word rates, whereas the TFS, the fast-varying component of speech, represents pitch, formant structure, timbre, etc. While envelope cues alone may be sufficient for partial speech comprehension in distortion free listening conditions, TFS is also important for speech comprehension, and especially so in the presence of distortions and competing backgrounds (Ding et al. 2013; Drullman 1995; Drullman et al. 1994a; 1994b; Kong et al. 2015; Moon and Hong 2014; Moore 2008; Rimmele et al. 2015; Smith et al. 2002; Swaminathan et al. 2016). Reverberation and noise affect the speech signal distinctly. While additive noise degrades the speech signal by reducing the intensity contrast, i.e., the depth of modulations, it does not affect the temporal sharpness of the speech signal. In contrast, reverberation, due to its convolutive nature, causes temporal smearing of both the envelope (example shown in Figure 2A, top) and TFS (see review by Assmann and Summerfield (2004)). TFS smearing results in spectral blurring (Figure 2A, bottom), which can affect the quality of the formant structure, timbre, and even pitch, whereas envelope smearing (Figure 2A, top) affects timing cues in the speech signal such as phoneme and syllable onset and offset. Such distortions can cause difficulties in identifying and discriminating consonants (Nabelek et al. 1989), vowels (Nabelek and Letowski 1988) and formant cues in reverberant listening conditions (Nabelek and Dagenais 1986).

**Figure 2.**
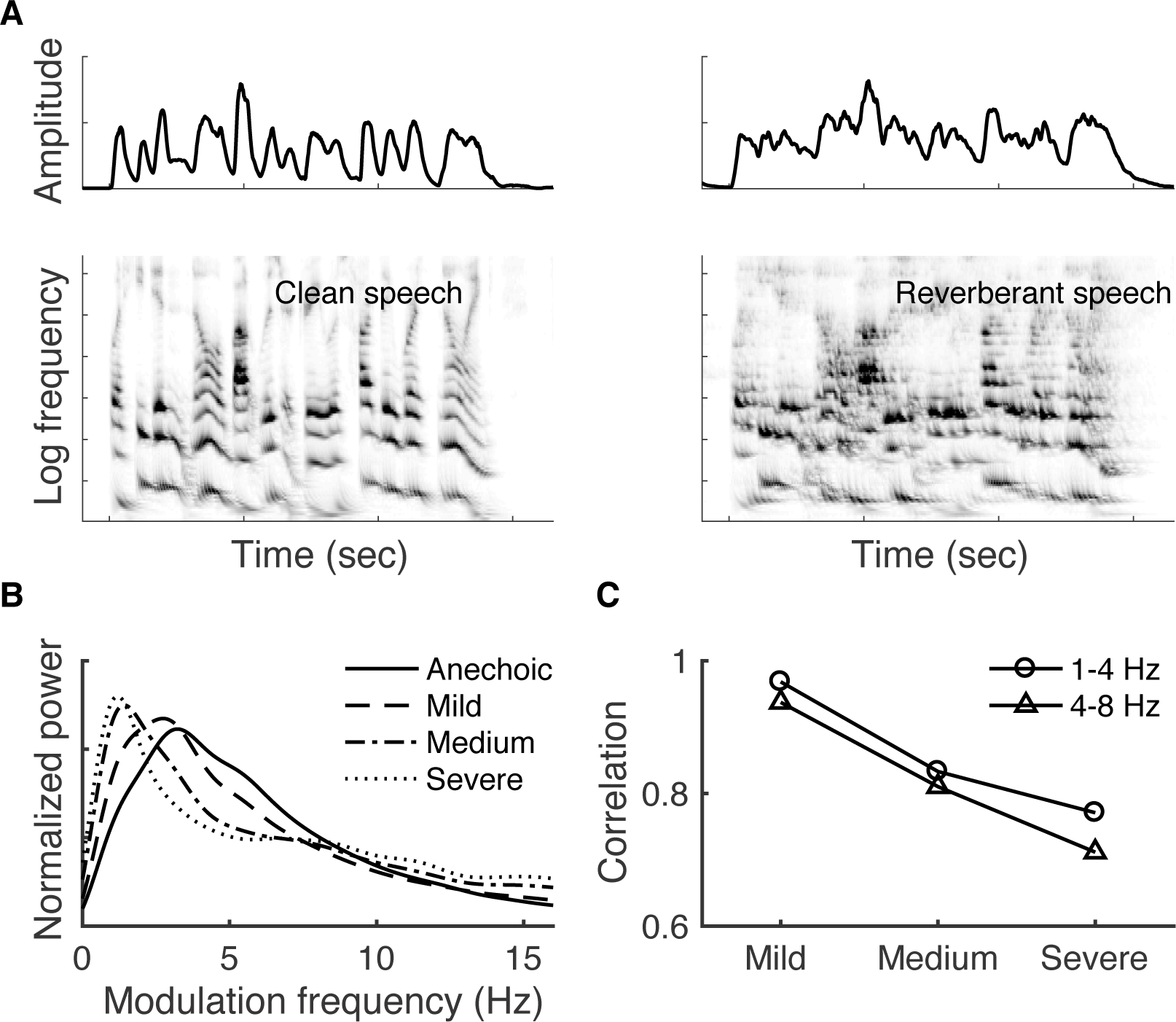
Effects of reverberation. **A.** Reverberation smears the temporal envelope (top right) of Clean speech (top left) as multiple reflections superimpose on the direct component from source. Reverberation also distorts the spectral structure of speech as shown by the auditory spectrogram (bottom) of speech without (left) and with (right) reverberation. **B.** The peak of the modulation spectrum occurs around 4 – 5 Hz in clean speech and shifts downward (left) with increasing severity of reverberation. **C**. Correlation coefficients comparing the bandpassed envelopes of reverberant speech, at different levels of severity, with the corresponding clean speech. The distortion effect of reverberation is higher in the 4 – 8 Hz band (corresponding to neural theta activity) than 1-4 Hz band (corresponding to neural delta activity).

Physiological studies, both in animal models (Mesgarani et al. 2014b; Moore et al. 2013; Rabinowitz et al. 2013) and humans (Ding and Simon 2013) have demonstrated the robustness of cortical representation of speech in the presence of stationary noise, in spite of degraded representation at the periphery of the auditory system (Delgutte 1980). Studies of the auditory brainstem (Fujihira et al. 2017; Sayles et al. 2014; Sayles and Winter 2008) and midbrain (Bidelman 2017; Devore and Delgutte 2010; Kuwada et al. 2014; Slama and Delgutte 2015) have shown that peripheral and subcortical neural coding of the temporal envelope can be substantially degraded in a reverberant environment. However, the effects of distortion due to reverberation, as well as the interaction of reverberation and additive noise (if any), on the cortical coding of speech, are less understood.

Here, using Magnetoencephalography (MEG) recordings of human subjects listening to continuous speech, and linear system methods of neural response prediction (encoding) and stimulus reconstruction (decoding) (Di Liberto et al. 2015; Ding and Simon 2012b; Pasley et al. 2012), we investigated the effect of noise and reverberation on cortical representation of continuous speech. Mesgarani et al. (2014b) examined the neural responses from single-unit recordings in ferrets, listening to reverberant speech (in absence of additive noise), and found that the corresponding clean speech spectrogram was better reconstructed than reverberant speech spectrogram. Further, Fuglsang et al. (2017), using electroencephalography (EEG) recordings of human subjects listening to speech in reverberation, showed that the clean speech envelope was better reconstructed than the reverberant speech envelope in case of severe reverberant listening conditions. The current study is formulated from a different point of view, (1) to systematically examine the joint effect of noise and reverberation on neural encoding of speech by varying the severity of both reverberation and noise, and (2) to examine the cortical representation of speech in noisy reverberant environment from both encoding and decoding perspectives, allowing insights into reverberation processing strategies across auditory cortex. We will show evidence that, while auditory cortex does show strong evidence of cleaning the speech signal of reverberation, the reverberant speech signal is also strongly represented, and many areas represent both reverberant and cleaned versions of the speech signal.

## Materials and Methods

### Subjects and Experimental Design

Thirteen normal-hearing, young adults participated in the experiment. All subjects were paid for their participation. The experimental procedures were approved by the University of Maryland Institutional Review Board and written informed consent was obtained from each subject before the experiment. Subjects listened to 60 s duration speech segments under a full factorial design of three noise and four reverberation levels, totaling twelve stimulus conditions. The three noise levels are No-noise, +6 dB and +3 dB signal-to-noise ratio (SNR). The four reverberation levels are referred to, with increasing severity, as anechoic (clean), mild, medium and severe reverberation with Reverberation Time to 20 dB (RT_20_: time elapsed before the reflections decay by 20 dB with respect to the direct component in terms of energy) values of 0 ms, 150 ms, 300 ms and 450 ms, respectively. The choice to use here the RT_20_ to characterize reverberation instead of the more standard RT_60_ (time elapsed before the reflections decay by 60 dB with respect to the direct component) arises from the usage of continuous reverberant speech: when speech reflections from an earlier time act as a masker for speech at the present time, a target-to-masker ratio (TMR) of 20 dB is perceptually more relevant than a TMR of 60 dB (which is instead more relevant for detection of reverberation in silence). In practice, any RT_20_ value is approximately one third of the corresponding RT_60_ value. Reverberant speech was generated by convolving a (base) clean speech segment with a Room Impulse Response (RIR) with the desired severity of reverberation. RIRs were generated using the image-source method (Allen and Berkley 1979) as implemented by Lehmann and Johansson (2010) by simulating listening conditions in a room of dimensions 7 x 5 x 3 m (length, width, height), with source and listener positioned at (4.5, 2.5, 1.7) m and (3, 2.5, 1.7) m, respectively. Different levels of reverberation were obtained by varying absorption coefficients of walls, floor and roof of the simulated room. Noisy reverberant speech was generated by adding spectrally matched noise to the reverberant speech, at the desired SNR; spectrally matched noise was generated by randomizing the phase of the reverberant speech signal and scaling it appropriately to achieve the required SNR. Mathematically, the stimulus *S* (*t*) is constructed as,

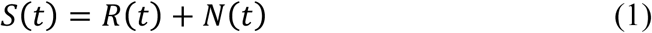

where *R*(*t*) and *N*(*t*) are respectively, the reverberant speech component of the stimulus and spectrally matched noise. *R*(*t*) is constructed as,

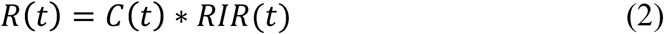

where *C*(*t*) and *RIR*(*t*) are, respectively, the (base) clean speech and the RIR. All twelve (base) speech segments, used to generate twelve stimulus conditions, were taken from a public domain narration of Grimms’ Fairy Tales by Jacob & Wilhelm Grimm (https://librivox.org/fairy-tales-by-the-brothers-grimm/), spoken by the same narrator. Periods of silence longer than 300 ms were replaced by a shorter gap whose duration was chosen randomly between 200 ms and 300 ms. To reduce loudness effects as a confound, when reverberation was added, the amplitude was rescaled so that all exemplars were of approximately equal perceptual loudness. No further scaling was performed when noise was added. Each of the twelve stimulus conditions was presented three times (trials) in succession, with the base speech segment used to generate a particular stimulus condition as well as presentation order of conditions randomized across subjects. To ensure the listeners’ attention, a target-word was set before each trial and the subjects were asked to count the number of occurrences of the target-word in the stimulus being played. Additionally, at the end of each trial, subjects answered a different 2-alternative-forced-choice comprehension question. Subjects were required to close their eyes while listening.

### Data recording and pre-processing

MEG recordings were conducted using a 160-channel whole-head system (Kanazawa Institute of Technology, Kanazawa, Japan). Its detection coils are arranged in a uniform array on a helmet-shaped surface of the bottom of the dewar, with ~25 mm between the centers of two adjacent 15.5-mm-diameter coils. Sensors are configured as first-order axial gradiometers with a baseline of 50 mm; their field sensitivities are 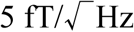 or better in the white noise region. Subjects lay horizontally in a dimly lit magnetically shielded room (Yokogawa Electric Corporation). Responses were recorded with a sampling rate of 1 kHz with an online 200-Hz low-pass filter and 60 Hz notch filter. Three reference magnetic sensors and three vibrational sensors were used to measure the environmental magnetic field and vibrations. The reference sensor recordings were utilized to reduce environmental noise from the MEG recordings using the Time-Shift PCA method (de Cheveigne and Simon 2007). Eye-blinks and heart beat artifacts were removed using Independent Component Analysis (ICA). For analysis in the sensor domain, MEG sensor recordings were decomposed into virtual sensors/components using denoising source separation (DSS) (de Cheveigne and Parra 2014; de Cheveigne and Simon 2008; Särelä and Valpola 2005), a blind source separation method that enhances neural activity consistent across trials. Specifically, DSS decomposes the multichannel MEG recording into temporally uncorrelated components, where each component is determined by maximizing its trial-to-trial reliability, measured by the correlation between the responses to the same stimulus in different trials. To reduce the computational complexity, sensor domain analysis was performed using DSS components. Additionally, for analysis in the source domain, each subject’s head shape was digitized (Polhemus 3SPACE FASTRAK) and the subject’s head was localized with respect to the MEG sensors using five marker coils attached to the head. The ‘fsaverage’ brain provided by FreeSurfer (Fischl 2012) was fit to each subject’s head shape using rotation, translation and uniform scaling. MEG data, after de-noising with time-shift PCA and ICA, were localized to active regions in the cortex using distributed minimum norm estimate (MNE) (Hamalainen and Ilmoniemi 1994) as implemented in MNE software (Gramfort et al. 2013; Gramfort et al. 2014). The source model comprised of 10242 regularly spaced virtual source dipoles in each hemisphere with orientations perpendicular to the cortical surface. The sensor noise covariance was estimated from the empty room recording data. Due to the auditory nature of the study, further analysis was restricted to the responses estimated at the sources located in the transverse, superior, middle temporal gyri and banks of the superior temporal sulcus (Desikan et al. 2006). Both speech envelope and neural response (either a DSS component in sensor space, or the estimated activity at one source domain location) were band pass filtered between 1 – 8 Hz (delta and theta bands), which correspond to the slow temporal modulations in speech (Ding and Simon 2012a; 2012b), for further analysis.

### Encoding of stimulus to neural responses

Encoding models provide a quantitative description of how information in a stimulus is represented in neural responses. Analyzing data from the perspective of encoding (predicting neural responses using the stimulus or some representation of the stimulus) allows investigators to identify, as well as quantify, how features/aspects of the stimulus are represented in the corresponding neural responses (Naselaris et al. 2011). Here, to identify the neural representation of speech distorted by noise and reverberation, three encoding models were compared namely: the Cleaned, Reverb and Mixed models (described below). Encoding analysis was performed by fitting a linear regression model between the stimulus representation under a particular model (whether Cleaned, Reverb or Mixed) and the corresponding low frequency (1-8 Hz) neural responses. This approach has been used previously to describe the temporal relation between a speech stimulus and the corresponding neural response as measured by MEG (Ding and Simon 2012b), EEG (Di Liberto et al. 2015), and ECoG (Mesgarani and Chang 2012). The resulting models are commonly referred to as Temporal Response Functions (TRFs) and are mathematically represented as

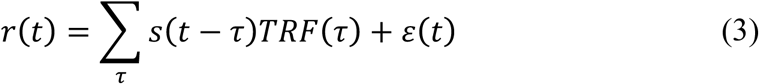

where *t* = 0,1,…, *T* are discretized time instances, *r*(*t*) is the neural response (of any individual sensor or DSS component, or the time-course of activity at a source location), *s*(*t*) is the choice of stimulus representation in the encoding model under consideration (referred to as ‘predictor’ here), *TRF*(*t*) is the TRF itself, and *∊*(*t*) is residual response waveform not explained by the TRF model (Ding and Simon 2012b). The TRF is estimated using boosting with 10-fold cross-validation (David et al. 2007). Success of the linear model, referred to as ‘prediction accuracy’, is evaluated by how well it predicts neural responses, as measured by the proportion of the variance explained: the square of the Pearson correlation coefficient between neural response *r*(*t*) and the TRF model prediction (right hand side of Eq. (3) excluding the error term). The three encoding models compared were: (1) the Cleaned model, where the stimulus is represented by the broadband envelope of the corresponding clean (base) speech, i.e. the envelope of *C*(*t*) of Eq. (2); (2) the Reverb model, where the stimulus is represented by the broadband envelope of the reverberant speech component of the stimulus, i.e. the envelope of *R*(*t*) of Eq. (1); and (3) the Mixed model – a model that allows both Cleaned and Reverb representations to contribute, i.e., simultaneously using envelopes from both the Cleaned and Reverb models as predictors. The Cleaned model tests the hypothesis that despite the distorted acoustic input to the ear, the cortex recovers and maintains neural representations for the underlying distortion free clean speech. The Reverb model tests the hypothesis that acoustic distortions due to reverberation present at the ear are also represented neurally in the cortex. Finally, the Mixed model allows the co-existence of neural representations for both clean and reverberant versions of speech. Such a dual representation is possible due to the hierarchical organization of the auditory cortex, which maintains increasingly complex and distortion robust representations of stimulus (Atencio et al. 2009; Okada et al. 2010; Sharpee et al. 2011). In all the encoding models, the broadband envelope was extracted by averaging the auditory spectrogram of the corresponding speech signal along the spectral dimension (Chi et al. 2005).

In case of the Mixed model, the linear model presented in (1) is modified as

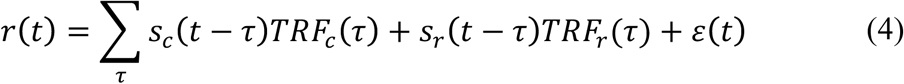

where *s_c_*(*t*) is the envelope of clean speech and *s_r_*(*t*) is the envelope of reverberant component of stimulus and *TRF_c_*(*t*),*TRF_r_*(*t*) are the corresponding TRFs. Due to the presence of two predictors, the Mixed model has twice the number of degrees of freedom than the corresponding Cleaned and Reverb models. To ensure that the increased accuracy (if any) of the Mixed model compared to the other two is merely not due to increased degrees of freedom, a non-informative speech envelope was added as an additional predictor in both Cleaned and Reverb models, thus balancing the number of free parameters across models. For example, in the Cleaned model, the non-informative speech envelope is obtained by replacing the first half of Reverb model envelope with its second half and vice versa.

### Decoding speech from neural responses

While the TRF/encoding analysis described in the previous section predicts neural response from stimulus, decoding analysis reconstructs stimulus envelope using neural responses. Thus, decoding analysis complements the TRF analysis (Mesgarani et al. 2009). Mathematically the envelope reconstruction/decoding operation is formulated as

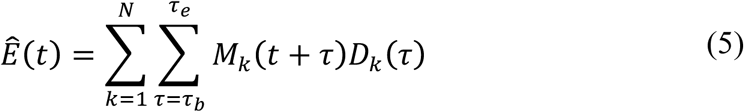

where 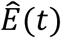 is the reconstructed envelope, *M_k_* (*t*) is the MEG recording (neural response) from sensor/component *k*, and *D_k_*(*t*) is the linear decoder for sensor/component *k*. The times *τ_b_* and *τ_e_* denote the beginning and end times of the integration window, 0 and 500 ms respectively here. The decoder is estimated using boosting, analogously to the TRF estimation in the previous section, to minimize the mean squared difference between reconstructed and actual envelopes. As decoding analysis linearly integrates information over all data (whether from all the sensors in sensor domain analysis or, equivalently, from all source points in source domain analysis) recorded in the time window under consideration, we restrict our decoding analysis to sensor space.

### Statistics

Due to the presence of multiple stimulus conditions (a total of 12 in the full factorial design with three noise and four reverberation levels), the following statistical approach was used to compare between different encoding or decoding models. Considering the example of comparison between Mixed and Reverb models, the difference between the two model prediction accuracies was calculated for each subject and condition and a repeated measures Analysis of Variance (ANOVA) is performed on the model differences with noise and reverberation as factors (Greenhouse-Geisser corrected when required). Significant effects were followed up with appropriate pairwise t-tests. Significant interaction effect was followed up with a t-test at each stimulus condition to compare the mean difference of models with zero, correcting for multiple comparisons using False Discovery Rate (FDR) (Benjamini and Hochberg 1995). In absence of a significant interaction effect, data was pooled according to the main effects, if present, before comparing the model differences against zero. Here also, FDR was used for multiple comparisons correction. For example, in the case of significant main effect for the reverberation factor but not noise, data was pooled across noise levels and a t-test was performed at each level of reverberation. When comparing two models, either in encoding or decoding analysis, through their differences, anechoic (reverberation free) stimuli were excluded as all models coincide in the anechoic listening condition and so differences would be identically equal to zero for all subjects, with zero variance.

In the case source domain analysis, nonparametric permutation tests (Maris and Oostenveld 2007; Nichols and Holmes 2002), based on the threshold-free cluster-enhancement algorithm (TFCE) (Smith and Nichols 2009), were used to control for multiple comparisons when testing for the significance of a result at a large number of source locations. The precise implementation details are available in the Eelbrain source code (Brodbeck 2017), but a brief summary follows. First, a test statistic (a t-value in case of t-test or an F-statistic in case of ANOVA) was computed for each source location based on the quantity of interest (here, the difference in prediction accuracies between two models) across subjects. The resulting test statistic map was then processed with TFCE, an image processing algorithm that enhances larger contiguous areas with large values compared to isolated spikes, based on the assumption that meaningful differences have a larger spatial extent than noise. To determine the null distribution for the resulting TFCE values, the procedure was repeated in 10,000 permutations of the data, with condition labels flipped for a randomly selected set of subjects in each permutation. The test statistic computation and TFCE were repeated in each permutation, and the largest value from the cluster-enhanced map is stored as an entry in the null distribution. Thus, a nonparametric distribution for the largest expected TFCE value under the null hypothesis was computed. Any value in the original TFCE map that exceeds the 95^th^ percentile of the distribution is thus significant at the 5% level. Thus, TFCE provides strong control over family-wise type-I error (Nichols and Holmes 2002).

## Results

To examine the neural representation of speech distorted by additive noise and reverberation, three possible encoding models were compared (Cleaned, Reverb and Mixed models; see Methods for detailed description) using neural responses from the first DSS (most dominant auditory) component (Ding and Simon 2012b). The performance of each model as measured by prediction accuracy (squared correlation coefficient between actual and predicted response) was computed for each model under each stimulus condition. In particular, if the brain maintains a distortion-free representation of speech in addition to the original distorted acoustic representation of speech, the Mixed model should have higher prediction accuracy than both the Reverb and Cleaned models, across all stimulus conditions. First, to compare the Mixed and Reverb models, repeated measures two-way ANOVA was performed on the difference of prediction accuracies between Mixed and Reverb models (Figure 3A) with noise and reverberation as within subject factors (anechoic level in reverberation factor was excluded as both models coincide when there is no reverberation). The main effect of reverberation was marginally significant (F(2, 24) = 3.307, p = 0.054) but no significant effect of noise was observed (F(2, 24) = 0.436, p = 0.652) along with no significant interaction (F(4, 48) = 0.112, p = 0.978). Post-hoc test at each reverberation level, after pooling data across noise levels, showed that model difference was significantly greater than zero at each reverberation level (mild: t(38) = 2.366, p = 0.023; medium: t(38) = 3.425, p = 0.002; severe: t(38) = 6.708, p < 0.001, multiple comparisons corrected via FDR with q = 0.05). This suggests that the Mixed model predicts neural responses better than the Reverb model across all stimulus conditions with reverberation. Similar comparison between Mixed and Cleaned models (Figure 3B) showed that model difference was significant both in noise (F(2, 24) = 14.380, p < 0.001) and reverberation (F(2, 24) = 13.546, p < 0.001) with significant interaction (F(4, 48) = 4.774, p = 0.003). Post-hoc tests at each stimulus condition showed that the model difference was significantly greater than zero at all reverberant conditions (FDR with q = 0.01). This suggests that the Mixed model predicts neural responses better than the Cleaned model. Taken together, these results suggest that when listening to speech in noisy reverberant conditions the auditory cortex maintains representations for both reverberant (distorted) and the corresponding clean (distortion free) versions of the stimulus.

**Figure 3.**
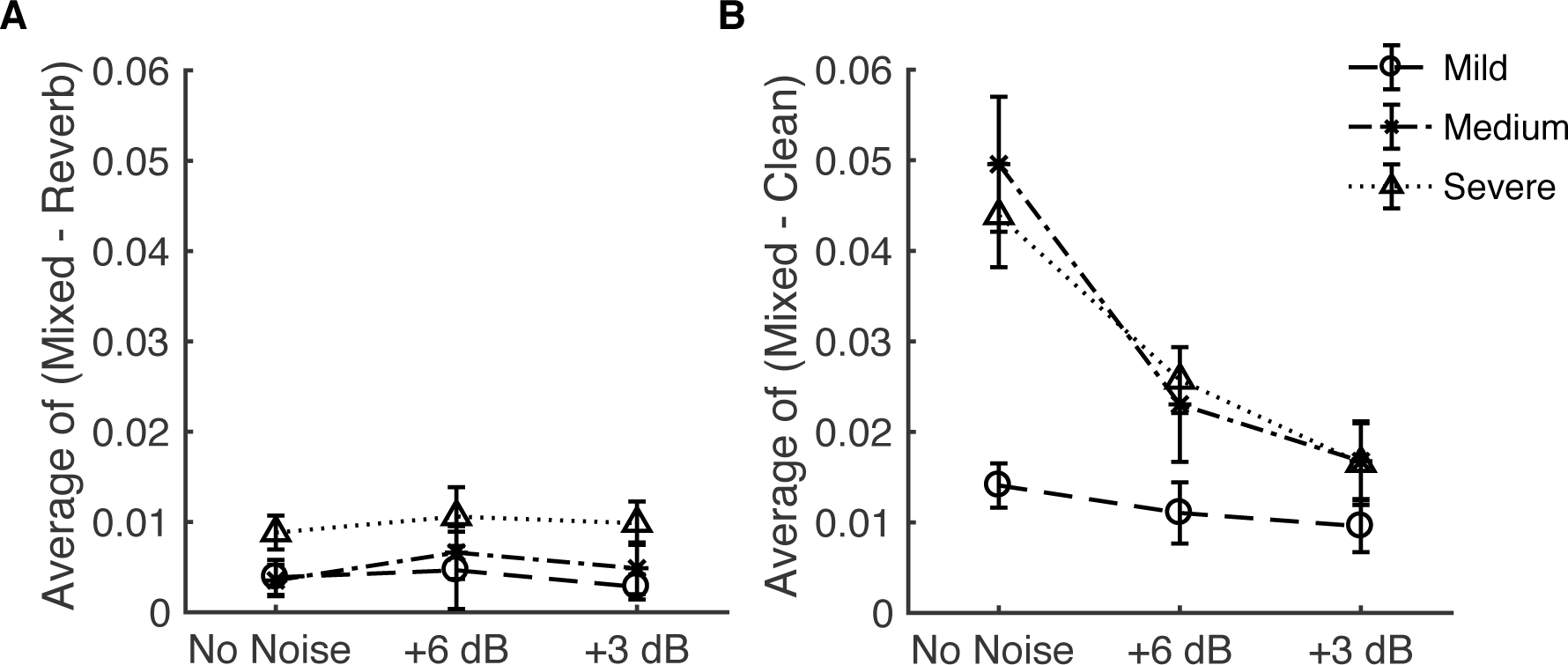
Comparing accuracy of encoding models. Difference between prediction accuracies of the Mixed and Reverb models (A) as well as Mixed and Cleaned models (B) are both significantly greater than zero (FDR at q = 0.05 and FDR at q =0.01 respectively). The Mixed model predicts neural responses significantly better than either the Reverb or Cleaned model for all stimulus conditions with reverberation.

To identify the cortical regions contributing to the increased prediction accuracy of the Mixed model compared with the Reverb model, encoding analysis was performed in the neural source domain (predicting neural activity at each source location). The difference between the prediction accuracies of the two models was computed at each source location for all stimulus conditions. Variation of model difference with respect to reverberation level was modeled, separately for each noise level, as the slope of a line fit between model difference and reverberation level, thus obtaining three data points (one value of slope per noise level) per source location. As ANOVA, correcting for multiple comparisons using TFCE, showed no significance with respect to noise (p >= 0.482), data was pooled by averaging the slope across three noise conditions, resulting in one value of slope per source location. Any value of slope significantly different from zero indicates significant model difference. A t-test performed at each source location, correcting for multiple comparisons, showed that Heschl’s gyrus and middle-to-posterior superior temporal gyrus areas contribute to the increased performance of the Mixed model over the Reverb model (Figure 4).

**Figure 4.**
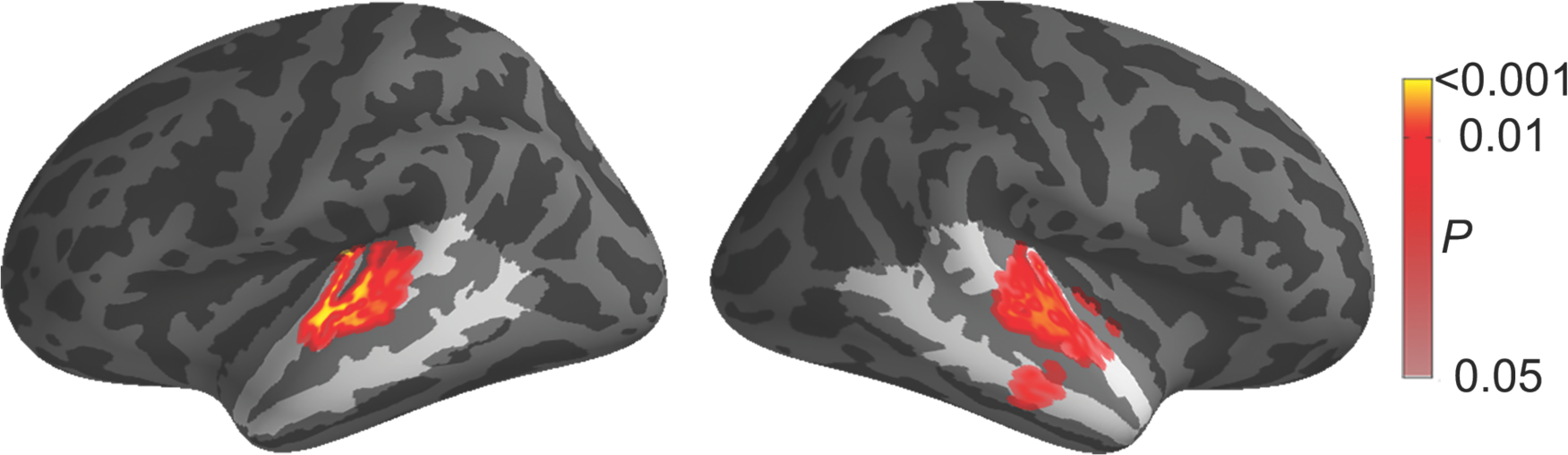
Anatomical regions contributing to increased performance of the Mixed model over the Reverb model (p < 0.05, corrected), rendered on the inflated brain surface model. These regions are better explained as containing areas with representations of both reverberant (distorted) and the corresponding clean (distortion free) versions of the stimulus, than as containing only representations of the reverberant (distorted) version. Areas that are not included in the analysis are shaded with a dark overlay.

To examine the fidelity of neural encoding of speech under different severity levels of noise and reverberation, prediction accuracies of the Mixed model (which best explained the neural response among the three encoding models compared) under different stimulus conditions were compared (Figure 5). A repeated measures ANOVA showed a significant interaction between noise and reverberation factors (F(2.761, 33.133) = 7.042, p = 0.001). Hence, post-hoc analysis was performed at each reverberation level to see the effect of noise. The variation of prediction accuracy with respect to noise, as measured by the slope of the line fit between noise levels and prediction accuracies, was calculated for each level of reverberation, per subject. A t-test at each reverberation level, corrected for multiple comparisons at q = 0.05 FDR, showed that the slope was significantly less than zero for mild (mean = −0.039, t(12) = −2.649, p = 0.021), medium (mean = −0.054, t(12) = −3.285, p = 0.007) and severe (mean = −0.047, t(12) = −3.410, p = 0.005) reverberation, whereas the anechoic condition showed no significant variation with respect to noise (mean = 0.0054, t(12) = 0.793, p = 0.443). This suggests that noise differentially affects the accuracy of neural encoding in listening conditions with and without reverberation: In the absence of any reverberation, noise did not show a significant effect on the accuracy of neural encoding, whereas its effect was adverse in presence of all reverberation levels tested. Because it is common in the experimental literature to not place too much emphasis on whether a speech stimulus is purely anechoic or instead contains mild reverberation (e.g. the case of a free field stimulus), it is worth re-emphasizing this result that anechoic listening condition has a representation that is markedly different from even the most mild reverberant listening condition. This can be seen in terms of their prediction accuracies when there is no noise (‘No Noise’ condition in Figure 5), as well as how the prediction accuracies change when noise is added.

**Figure 5.**
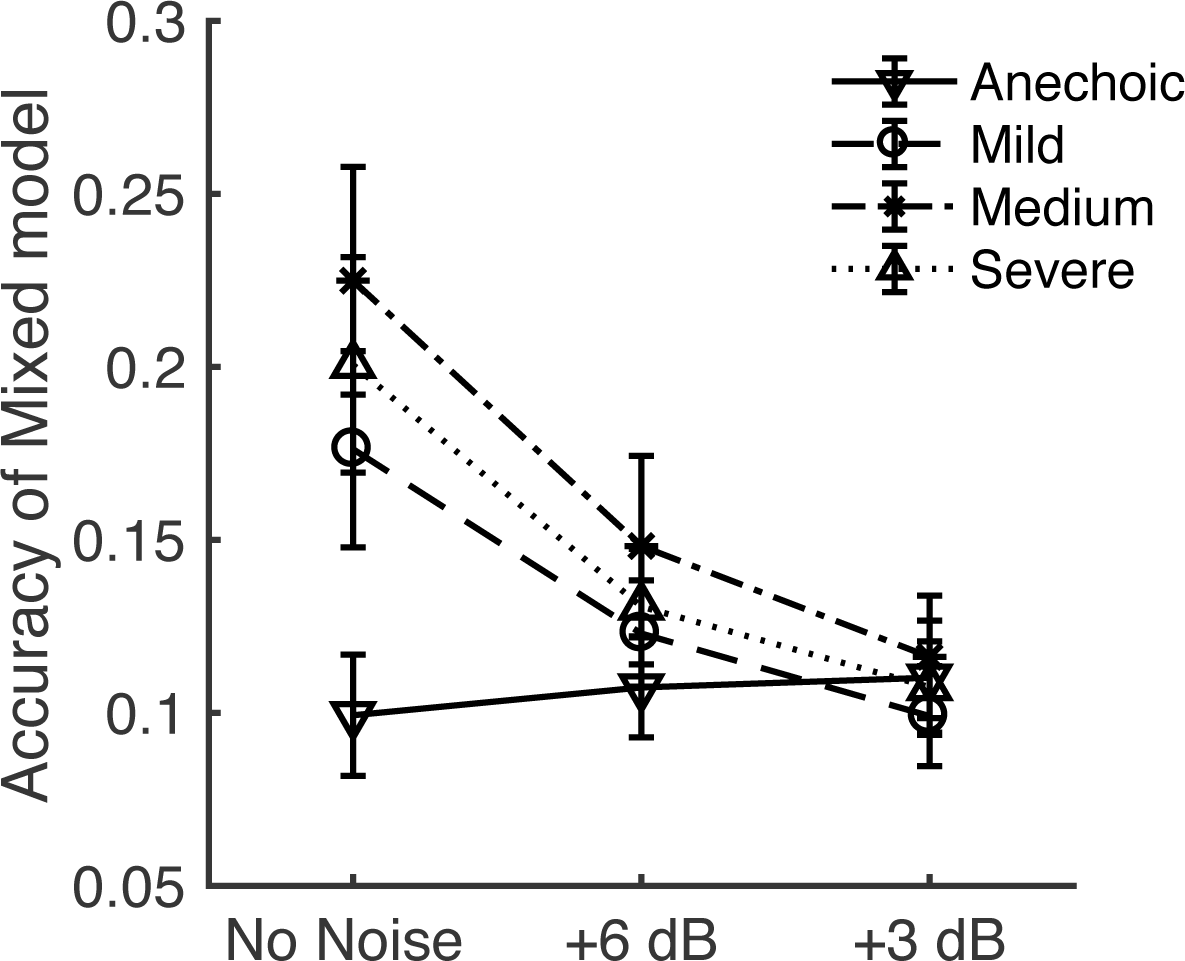
Effect of noise and reverberation on accuracy of neural encoding. In the absence of reverberation (“Anechoic”), noise did not show any significant effect on the accuracy of neural encoding. In contrast, encoding accuracy was reduced significantly with increase in noise, in the presence of *any* reverberation.

While the results presented so far provide an encoding perspective of speech in noisy and reverberant listening conditions, the putative role of delta and theta band neural responses in representing different aspects of speech (Ding and Simon 2014; Kösem and Van Wassenhove 2017) is examined in the following. The results from encoding models suggest that the auditory cortex maintains representations for both reverberant and cleaned versions of speech in reverberant environments. To assess the relative contributions of delta and theta band neural responses to the reverberant and cleaned representations, decoding analysis was employed. Here, both the reverberant and the respective clean versions of the stimulus envelope were reconstructed using delta and theta band neural responses separately, in order to compare which version of the envelope is more faithfully represented by delta and theta neural response. Figure 6 shows the difference between reconstruction accuracies of the reverberant and cleaned envelopes using only delta or only theta band neural responses. A repeated measures ANOVA of model differences (Reverb - Cleaned), in the delta band, showed a significant effect of noise (F(2, 24) = 7.005, p = 0.004), reverberation (F(2, 24) = 8.564, p = 0.002) as well as significant interaction (F(4, 48) = 3.019, p = 0.027). Post-hoc t-tests showed that model difference was significantly greater than zero in all reverberant stimulus conditions (multiple comparisons corrected via FDR at q = 0.05). Similar analysis using theta band neural responses showed that model differences were not significantly affected by noise (F(1.395, 16.743) = 0.265, p = 0.691) or reverberation (F(2, 24) = 0.904, p = 0.418) with no significant interaction effect (F(2.622, 31.463) = 2.034, p = 0.104). Further, post-hoc analysis showed that the model difference at any stimulus condition was not significantly different from zero (correcting for multiple comparisons using FDR). These results suggest that the delta band responses dominantly maintain reverberant representation, whereas theta band contains nearly equal contributions from both cleaned and reverberant representations. Delta band neural responses maintain a better representation of reverberant speech than cleaned, while theta band showed no such distinction.

**Figure 6.**
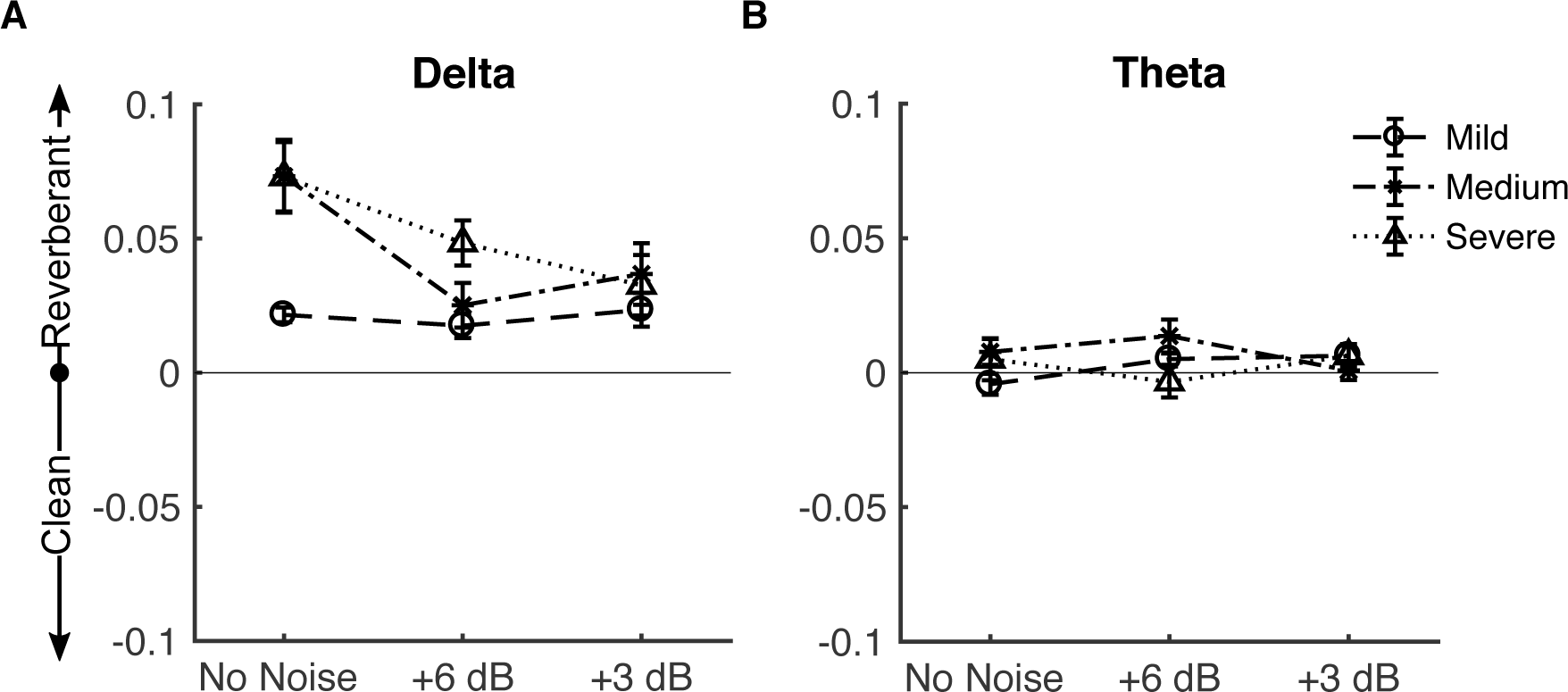
Comparing stimulus reconstruction accuracies for reverberant and corresponding clean speech. Results above the midline favor the Reverb model; below the midline favor the Cleaned model. **A.** Using only delta band (1 – 4 Hz) neural responses, the stimulus reconstruction of reverberant speech is significantly better than the corresponding clean speech (FDR with q = 0.05). **B.** Reconstruction using only theta band (4 – 8 Hz) neural responses did not show significant differences (FDR with q = 0.05) between reconstruction accuracies of the reverberant and respective clean stimulus.

## Discussion

Using MEG to record the cortical activity of subjects listening to noisy, reverberant speech, and linear methods of neural response prediction and stimulus reconstruction, we observed that (1) the cortex maintains both distorted as well as the corresponding distortion-free (cleaned) representations of speech (2) noise differentially affects the accuracy of neural encoding in absence and presence of reverberation (3) theta band neural responses are a more likely candidate than delta band neural responses to hold the distortion free representation of the (distorted) acoustic stimulus.

That the Mixed model has better encoding accuracy compared to both the Reverb and Cleaned models (Figure 3) suggests that both distorted (reverberant) and distortion free (cleaned) versions of the speech are represented in auditory cortex. Such a dual representation is feasible given the hierarchical nature of auditory processing in cortex (Okada et al. 2010), where progressively distortion free (Moore et al. 2013; Rabinowitz et al. 2013) and ultimately categorical representations of speech emerge (Chang et al. 2010; Di Liberto et al. 2015; Peelle et al. 2010). Reverberation cleaning is often tied to the phenomenon of echo suppression, typically investigated in simple stimuli such as lead-lag pairs where it is referred to as the precedence effect and is often explained using inhibition triggered by the leading sound (Litovsky et al. 1999; Xia and Shinn-Cunningham 2011). Mesgarani et al. (2014b) suggest a similar mechanistic model based on feed-forward synaptic depression and feed-back gain normalization to reduce the distortion due to reverberation. Traer and McDermott (2016) suggest that the problem of speech comprehension in reverberant conditions is solved by the auditory system as part of the general cocktail party problem due to its ill-posed nature. They suggest that the brain uses prior information, accumulated through experience, to separate the clean speech from distorted reverberant speech input to the ear and identify it as an auditory object, separate from the environment in which it was produced. Both of these approaches ((Mesgarani et al. 2014b) and (Traer and McDermott 2016)) argue for simultaneous cortical representations of cleaned speech and the original reverberant speech, as shown in the current study. Significant difference between the prediction accuracies of the Mixed and Reverb models, reflecting the contribution of the distortion free part of the Mixed model, was confined to Heschl’s gyrus and middle to posterior superior temporal gyrus (Figure 4). Similar anatomical areas have been implicated as the substrate of categorical (phonemic) representation of speech in the cortex (Mesgarani et al. 2014a), suggesting that the cleaned contribution of the Mixed model could be related to the computation of distortion invariant categorical representation of speech.

In the absence of reverberation, the accuracy of neural encoding of speech is not significantly affected by noise (Figure 5). Such robustness to stationary noise has been previously demonstrated (Ding and Simon 2013) and is thought to be the result of neural adaptation to statistics (such as mean and variance) of sound intensity (Dean et al. 2005; Dean et al. 2008; Robinson and McAlpine 2009). However, in the case of reverberant environments, our results show that the neural encoding of speech is strongly and detrimentally affected by the addition of stationary noise (Figure 5). A similar detrimental effect of stationary noise has been previously observed using vocoded speech (Ding et al. 2013), highlighting the importance of TFS integrity for accurate neural encoding of speech in noisy background in contrast to the quite listening conditions, wherein envelope cues are thought to be sufficient. Further, this suggests that the envelope entrainment to speech observed in MEG and EEG studies is a function of TFS along with the envelope.

On the other hand, in the absence of noise the encoding accuracy of reverberant speech (even under mild reverberation) is significantly higher compared with anechoic condition (Figure 5). The low-pass nature of the cortical response modulation transfer function (Simon and Ding 2010), combined with the downward shift of modulation spectrum with increasing reverberation (Figure 2B), could explain the increase in accuracy of neural encoding with reverberation in the absence of noise. However, the effect of listening effort due to reverberation cannot be discounted here either. Thus, the observed increase in encoding accuracy with increase in reverberation, in the absence of noise, could be due to combined effect of a change in modulation spectrum and listeners’ effort. Another distinct possibility could be due to the fact that reverberant listening conditions, even mild, are pervasive in daily life, whereas anechoic listening conditions are rarely experienced. Slama and Delgutte (2015), using an animal model, observed enhanced coding of amplitude modulated stimuli in reverberant environments compared with the anechoic condition. Thus, it is possible that ecologically irrelevant anechoic speech is not encoded as accurately as speech in ecologically relevant listening conditions.

Along with successful comprehension of speech in typical reverberant environments, a listener can also perceive and make subjective judgments regarding the reverberant environment, suggesting that such information is readily accessible to the auditory system. The observation that a reverberant envelope is better reconstructed than the corresponding cleaned envelope using only delta band neural responses (Figure 6A) suggest that the delta band is a candidate to convey the perception of reverberation. Similar reconstruction results using theta band neural responses (Figure 6B) showed no preference for either the reverberant or cleaned envelope. Despite the increased stimulus contrast (reduced correlation) between the reverberant and clean envelopes in the theta band compared to delta band (Figure 2C), the shift away from the reverberation-dominated decoding in delta to the more balanced representation in theta provides limited evidence for reverberation removal occurring dominantly in theta band neural responses. These observations are consistent with the hypothesized roles of slow varying delta band and fast varying theta band neural responses to encode information related to the perceived non-speech specific acoustic rhythm and speech specific modulations necessary for intelligibility respectively (Ding and Simon 2014). As such, it is beneficial for the auditory system to reduce the distortion in the theta band more than the delta band (Figure 6). In contrast to the decoding results presented here, using a combination of both delta and theta band neural responses, Fuglsang et al. (2017) showed that cleaned speech envelope was better reconstructed than reverberant speech envelope in case of severe reverberation. This difference may be due to the lack of binaural cues in the current study, which are known to enhance speech perception in reverberant and noisy environments (Nabelek and Robinson 1982). Also, using single unit recording from the primary auditory cortex of ferrets, Mesgarani et al. (2014b) showed that cleaned speech was better reconstructed when listening in reverberant conditions. This difference with the decoding results presented here could be due to the availability of spike/high-gamma (> 40 Hz) neuronal responses in single unit recordings, in contrast to the current study, which examined only slow temporal modulations.

In summary, the results suggest that while listening to speech distorted by additive noise and reverberation, the auditory cortex maintains representations for both distorted and the corresponding cleaned (distortion free) speech. The additive noise differentially affects the accuracy of neural encoding in presence and absence of reverberation. Finally, theta band neural responses are a candidate for containing distortion free representations of speech in reverberant environments, while the delta band neural responses may convey the non-speech-specific information regarding the reverberant listening environment.

## Author Contributions

M.V.D conceived the study; K.C.P, M.V.D and J.Z.S designed the study; K.C.P performed the experiment; K.C.P, C.B and J.Z.S analyzed the data; K.C.P and J.Z.S prepared the manuscript.

## Acknowledgements

The authors thank Natalia Lapinskaya for excellent MEG technical support.

